# Soil chloride content influences the response of bacterial but not fungal diversity to silver nanoparticles entering soil via wastewater treatment processing

**DOI:** 10.1101/715839

**Authors:** Christian Forstner, Thomas G. Orton, Peng Wang, Peter M. Kopittke, Paul G. Dennis

**Affiliations:** School of Earth and Environmental Sciences, The University of Queensland, St Lucia, Brisbane, QLD 4072, Australia; School of Agriculture and Food Sciences, The University of Queensland, St Lucia, Brisbane, QLD 4072, Australia

**Author notes:** Address correspondence to Paul G. Dennis. College of Resources and Environmental Sciences, Nanjing Agricultural University, Nanjing, China (Peng Wang).

**Keywords:** phylogenetic marker gene sequencing, nanoparticles, microbial diversity, nanotechnology, environmental risk

## Abstract

Silver nanoparticles (NPs) are among the most widely used nanomaterials and are entering soil ecosystems, mainly via biosolids in agriculture. When added directly to soils, metallic Ag-NPs have been shown to affect microbial communities, which underpin important ecosystem functions. During wastewater treatment processing, metallic Ag-NPs are rapidly converted to Ag_2_S, which is relatively insoluble and less toxic. Furthermore, recent evidence indicates that silver bioavailability is influenced by soil chloride content. Hence there is a need to understand the impacts of wastewater treatment processed Ag-NPs at varying levels of salinity on soil microbial diversity. In this study, we examined how the application of 0 g, 1 g and 2 g kg^−1^ NaCl to soil influence the effects of 0 mg, 1 mg and 10 mg kg^−1^ Ag, applied as wastewater treatment processed Ag-NPs, on bacterial and fungal diversity over time. Using high-throughput phylogenetic marker gene sequencing we demonstrate that, despite being theoretically less toxic, wastewater treatment processed Ag-NPs can affect the composition of soil bacterial and fungal communities, and influence bacterial alpha diversity. In addition, we found that silver-associated changes in bacterial community composition were affected by soil chloride content, with more acute responses to silver being observed in more saline soils. This work highlights that the release of Ag-NPs into soils via realistic exposure pathways can alter microbial diversity and that these effects may be influenced by soil chloride content.

**Summary capsule:** Soil chloride content influences the response of bacterial but not fungal diversity to wastewater treatment processed silver nanoparticles.

## 1. Introduction

Silver nanoparticles (Ag-NPs) are used for a wide-range of applications (Vance et al., 2015) and are projected to have a total market value of US$3 billion by 2021 (Inshakova and Inshakov, 2017). As a result of their widespread use, they are also being released into the broader environment, mainly via biosolids in agriculture (Gottschalk et al., 2009; Sun et al., 2016, 2015; Voelker et al., 2015; Wang et al., 2018b). The addition of freshly synthesized metallic Ag-NPs to soil has been shown to influence bacterial diversity (Samarajeewa et al., 2017). In wastewater treatment systems and soils, however, metallic Ag-NPs are typically converted to silver sulfide nanoparticles (Ag_2_S-NPs), which are less toxic to microbes and other organisms (Judy et al., 2015; Levard et al., 2013; Reinsch et al., 2012; Wang et al., 2018a). Hence, the impacts on soil microbial diversity of metallic Ag-NPs that have been processed and applied in an environmentally realistic manner are poorly understood. In addition, soil chloride content, has been shown to enhance the bioavailability of silver in soils in which metallic Ag-NPs had been largely converted to Ag_2_S during the wastewater treatment process (Wang et al., 2016). At present, however, it is not known whether soil chloride concentration influences the impacts of environmentally relevant forms of Ag-NPs (i.e. Ag_2_S) on microbial diversity. This information is important for the safe and sustainable use of nanomaterials.

Of all silver entering wastewater treatment systems, up to 70% is applied to agricultural land in biosolids (Kaegi et al., 2011; Lovingood et al., 2018), in which 85-100% is present as Ag_2_S (Doolette et al., 2013; Kaegi et al., 2015, 2011; Lombi et al., 2013; Ma et al., 2014; Pradas Del Real et al., 2016; Wang et al., 2018a, 2016). This is critical, because the solubility of Ag_2_S (Ksp = 6 × 10^−51^) is markedly lower than for metallic Ag-NPs, and hence Ag_2_S is presumably less toxic to soil organisms. Despite this, most studies concerning the effects of Ag-NPs on soil microbial communities have focused on freshly synthesized metallic Ag-NPs, which have been shown to decrease microbial diversity (Samarajeewa et al., 2017), biomass (Grün et al., 2018) and enzyme activity (Eivazi et al., 2018; Grün et al., 2018; Samarajeewa et al., 2017), as well as the abundances of nitrogen-fixing organisms (Grün et al., 2018; Kumar et al., 2011). Relative to freshly synthesized metallic Ag-NPs and ionic silver, Judy et al. (2015) found that Ag_2_S-NPs led to smaller reductions in microbial biomass, were less disruptive to plant-mycorrhizal symbioses and were less phytotoxic. The reduced toxicity of Ag_2_S-NPs is likely due to the occlusion of bioavailable Ag^+^ by the insoluble Ag_2_S shell that forms on the surfaces of Ag-NPs as they undergo sulfidation (Elechiguerra et al., 2005; Kaegi et al., 2011; Levard et al., 2013). This phenomenon may also explain why Judy et al. (2015) found that the effects of Ag_2_S on microbial communities are not dose dependent. In addition to changes in the speciation of silver during wastewater treatment, nanoparticles become integrated into the biosolids (Schlich et al., 2018). Hence when contaminated biosolids are applied to agricultural land, silver may encounter soil microbes less frequently than if non-contaminated biosolids and metallic Ag-NPs were added to soil separately. This is the case in most studies and may exaggerate the impacts of wastewater treatment processed Ag-NPs on soil microbial communities.

Once in soil, the toxicity of silver from wastewater treatment pathways, is further modified by the properties of the soil itself (Rahmatpour et al., 2017; Schlich and Hund-Rinke, 2015). Previously, using plant bioassays and diffusive gradients in thin films, we have demonstrated that soil chloride content is positively associated with the bioavailability of silver despite the metallic Ag-NPs having been converted largely to Ag_2_S in the sludge (Wang et al., 2016). Hence, soil microorganisms are also likely to be exposed to elevated levels of toxic Ag^+^ ions when biosolids are added to saline soils. Furthermore, microbial communities in saline soils are typically less diverse than those in non-saline soils (Zhang et al., 2019) and may be more vulnerable to perturbation. Despite this, the influence of chloride content on the impacts of wastewater treatment processed Ag-NPs on soil microbial diversity are not known.

In this study, we examined how the application of 0 g, 1 g and 2 g kg^−1^ NaCl to soil influences the effects of 0 mg, 1 mg and 10 mg kg^−1^ Ag, applied as wastewater treatment processed Ag-NPs, on bacterial and fungal diversity. These NaCl doses resulted in soil solution electrical conductivities (EC) of 1, 3 and 6 dS m^−1^, which assuming a threshold of 4 dS m^−1^ for a saline soil (United States Salinity Laboaratory Staff, 1954), represented a non-saline, sub-saline and saline soil, respectively. Metallic Ag-NPs and wastewater were processed for 28 days in sequencing batch reactors to generate sludge that was applied to soils to achieve silver concentrations equivalent to the 95^th^ percentile of one and ten years of release according to EPA surveys (United States Environmental Protection Agency, 2009; Wang et al., 2016).

Sludges were applied at a rate of 870 g m^−2^ soil, mimicking EPA guidelines (United States Environmental Protection Agency, 2009; Wang et al., 2016). A control sludge without added Ag was applied to control soils at the same rate. All treatments were replicated three times and communities were characterized at 3, 7, 30, and 90 days post-inoculation using high throughput phylogenetic marker gene sequencing. We hypothesized that: 1) Ag-NPs entering soil via wastewater treatment processing would influence the diversity of bacterial and fungal communities, and that 2) these effects would be more apparent in soils with elevated chloride content, as chloride has been shown to increase the bioavailability of silver.

## 2. Materials and methods

### 2.1. Experimental design

The soil used in this study is classified as a Kandosol according to the Australian Soil Classification (Isbell, 2002), or alternatively classed as an Ultisol according to the USDA Soil Taxonomy (Soil Survey Staff, 2014) and has been described in our previous work (see Table S1 from Wang et al. (2016)). Briefly, the soil was collected at a depth of 0−20 cm from a pineapple (*Ananas comosus*) farm in Queensland, Australia (27.02 °S, 152.92 °E). The soil had a sandy loam texture, pH of 5.4 (1:5 soil/water), soil solution EC of 1.0 dS/m (saturation extract), a cation exchange capacity of 2.6 cmol_c_/kg and a total organic C content of 1.1% (Wang et al., 2016). Fresh soil was passed through a 2 mm sieve and treated with 0, 1 or 2 g kg^−1^ NaCl in order to investigate the effects of chloride addition. This increased the EC to 3 dS m^−1^ and 6 dS m^−1^ respectively. Following the addition of NaCl, deionized water was added until 60% field capacity was attained, and soils were left to equilibrate for one month. Soils were then split into nine c. 1.6 kg sub-samples to which the silver treatments were applied

### 2.2. Sludge generation

In order to simulate a realistic exposure pathway of Ag-NP application to soils, two sludges were generated in sequencing batch reactors with working volumes of 10 L and initial mixed liquor suspended soil concentration of 4 ± 0.2 g L^−1^. Reactors were prepared according to Doolette et al. (2013) and details have been previously given in Wang et al. (2016). Briefly, one reactor was amended with metallic Ag-NPs (equiv. to OECD NM300, Fraunhofer IEM, 15-20 nm) while the other served as a control. Each feed cycle lasted 24 h with amendments being added at the beginning of each cycle. After 28 days of reactor operation, sludges were collected for subsequent use. In order to achieve realistic Ag-NP concentrations within the treatment sludges, the Ag-NP sludge was mixed with control sludge to achieve final Ag concentrations of 57 and 570 mg kg^−1^. These concentrations conform to the 95^th^ confidence interval as well as a ten-fold increase of Ag concentrations in sludges of publicly owned wastewater treatment works in the U.S.A. (United States Environmental Protection Agency, 2009; Wang et al., 2016). Using this experimental system, we have demonstrated that 87% of the metallic Ag-NPs initially added to the sludge are converted to Ag_2_S, with this Ag_2_S stable in both the sludge and the soil upon its subsequent addition (Wang et al., 2016).

### 2.3. Sludge application to soils

The mass of sludge applied to each soil was equivalent to U.S. EPA guidelines (870 g m^−2^) (Stein et al., 1995), calculated as 17.5 g kg^−1^ (assuming 8 cm depth of incorporation of soil with a density of 1.25 cm^−3^ and biannual applications) (Colorado State University, 1995). The application of Ag-containing sludges resulted in silver concentrations of 1 and 10 mg Ag kg^−1^ soil. The control sludge contained a low level of background silver, resulting in a final concentration of 0.02 mg kg^−1^ in soils. Three replicate 500 g samples of each treatment were placed into 1 L plastic containers with lids that facilitated gas exchange. This yielded 27 containers that were incubated for 90 days in the dark at 25°C. Humidity was maintained at 80% in order to keep the soils at the same WHC throughout the experimental period.

### 2.4. Soil sampling and DNA extraction

Soil cores, (~25 g) were collected from each experimental unit after 3, 7, 30 and 90 days using sterile 50 ml plastic tubes, and immediately transferred to −80°C storage. DNA extraction was conducted on 250 mg of thawed soil using the Power Soil DNA Isolation kit (MO BIO Laboratories, Carlsbad, CA) according to the manufacturers’ instructions.

### 2.5. Bacterial 16S rRNA and fungal ITS2 gene amplification, and sequencing

Universal bacterial 16S rRNA genes were amplified by polymerase chain reaction (PCR) and subsequently processed according to Forstner et al. (2019). Briefly, bacterial 16S rRNA genes were PCR amplified using 926F (Engelbrektson et al., 2010) and 1392wR (Engelbrektson et al., 2010) primers and sequencing was conducted on an Illumina MiSeq using a MiSeq Reagent Kit v3 (600 cycles; Illumina) and 30% PhiX Control v3 (Illumina).

Fungal ITS2 regions were PCR amplified using the primers gITS7F (5’- GTG ART CAT CGA RTC TTT G -3’) (Ihrmark et al., 2012) and ITS4R (5’- TCC TCC GCT TAT TGA TAT GC -3’) (White et al., 1990). PCRs were performed on 2.5 μl DNA samples in 1× Phire Green Reaction Buffer (Thermo Fisher), 100 μM of each of the dNTPs (Invitrogen), 0.5 μl Phire Green Hot Start II DNA Polymerase, and 250 μM of each primer, made up to a total volume of 25 μl with molecular biology grade water. Thermocycling conditions were as follows: 98°C for 45 seconds; then 35 cycles of 98°C for 5 seconds, 55°C for 5 seconds, 72°C for 6 seconds; followed by 72°C for 1 minute. Amplifications were performed using a Veriti^®^ 96-well Thermocycler (Applied Biosystems). PCR success was determined by gel electrophoresis, which also facilitated visual confirmation of amplicon size and quality. Subsequent processing and sequencing of amplicons on an Illumina MiSeq was conducted as described above for 16S rRNA gene amplicons as well as in Forstner et al. (2019).

### 2.6. Processing of sequence data

Data were analyzed as described in Forstner et al. (2019). Briefly, both 16S and ITS2 datasets were analyzed using the forward reads only. For 16S rDNA sequences, USEARCH (v10.0.240) (Edgar, 2010) was used for primer removal and trimming to 250 bp using fastx_truncate. High-quality sequences were then identified using fastq_filter (−fastq_maxee=1) and duplicate sequences were removed using fastx_uniques. Sequences were clustered at 97% similarity into operational taxonomic units (OTU) and potential chimeras were identified and removed (cluster_otus). Lastly an OTU table was generated using otutab with default parameters from the pre-trimmed reads and the OTU representative sequences.

For the ITS data, ITSx v1.0.11 (Bengtsson-Palme et al., 2013) was used to identify and extract fungal ITS2 sequences. Chimeric ITS2 sequences were identified and removed using the uchime2_ref command of USEARCH and the UNITE database (v7.2 - 2017.10.10) (Nilsson et al., 2019). After this, the sequences were then clustered at 97% similarity into operational taxonomic units (OTU) and an OTU table was generated using the otutab command of USEARCH with default parameters. SILVA SSU (v128) (Quast et al., 2013) and UNITE (v7.2-2017.10.10) (Nilsson et al., 2019) taxonomy was assigned to the 16S and ITS sequences, respectively, using BLASTN (v2.3.0+) (Zhang et al., 2000) within the feature classifier of QIIME2 (v2017.9) (Boylen et al., 2018). The 16S OTU table was then filtered to remove OTUs classified as chloroplasts, mitochondria, archaea or eukaryotes using the BIOM (McDonald et al., 2012) tool suite. For the 16S sequences, de-novo multiple sequence alignments of the representative OTU sequences were generated using MAFFT (v7.221) (Katoh and Standley, 2013) and masked with alignment mask (QIIME2). This masked alignment was used to generate a midpoint-rooted phylogenetic tree using FastTree (v2.1.9) (Price et al., 2010) in QIIME2. OTU tables were rarefied to 4850 and 1150 sequences per sample for 16S and ITS, respectively. The mean numbers of observed (Sobs) and predicted (Chao1) OTUs as well as Shannons’ diversity index were calculated using QIIME2 for both bacteria and fungi.

### 2.7. Statistical analyses

For statistical analyses, we considered the effects of the applied dose of Ag-NPs (0, 1 or 10 mg kg^−1^ soil), the applied dose of chloride (0, 1 g or 2 g kg^−1^ soil), and day (3, 7, 30, and 90 days post inoculation); here we refer to these factors as: *Ag-NP*, *Cl* and *Day*. To determine whether the main and interaction effects of *Ag-NP*, *Cl* and *Day* significantly affected alpha diversity and the relative abundances of individual taxa we used a linear mixed-effects modeling approach (Pinheiro and Bates, 2004). In these models *Ag-NP*, *Cl* and *Day*, as well as their two-way and three-way interactions, were treated as fixed effects, and soil containers (samples) were treated as a random effect to account for the repeated measures. F-tests were applied to assess significance (*P* < 0.05), and were implemented in R using the *lme4* (Bates et al., 2015) and *lmerTest* (Kuznetsova, 2017) packages.

Differences in the composition of microbial communities between samples (beta diversity) were assessed using multivariate generalized linear models using a negative binomial distribution (Warton, 2011). The significance of differences in community composition was determined by comparing the sum-of-likelihood test statistics for the alternative statistical models via a resampling method (Wang et al., 2012) that accounted for the correlation between species and the correlation within the repeated measures taken from the same sample container. These comparisons were implemented in R using the *mvabund* package (Wang et al., 2012). Taxa whose maximum relative abundance was less than 0.1% for bacteria or 0.4% for fungi were disregarded before statistical analysis. Post-hoc analyses were undertaken to investigate which doses of Ag-NPs and chloride differed on what days. The Benjamini-Hochberg correction was applied to all post-hoc tests.

### 2.8. Data availability

All sequences have been deposited to the Sequence Read Archive (SRA) under BioProject accession number PRJNA543161.

## 3. Results

### 3.1. Soil bacterial community composition

Bacterial communities were dominated by representatives of the Acidobacteria, Actinobacteria, Armatimonadetes, Bacteroidetes, Chloroflexi, Cyanobacteria, Fibrobacteres, Firmicutes, Gemmatimonadetes, Microgenomates, Nitrospirae, Planctomycetes, Proteobacteria, Saccharibacteria, Spirochaetae and Verrucomicrobia (Fig. S1).

Multivariate GLMs indicated that bacterial community composition was significantly influenced by silver, chloride, time and their interactions (Tables 1 and 2). The magnitude of compositional change in response to silver diminished faster at higher chloride levels, with differences at 90 days being considerably smaller that at 3, 7 and 30 days (Fig. 1).

**Table 1.** *P*-values from multivariate tests computed using mvabund highlighting the main and interaction effects of silver, chloride and day on microbial community composition. Significant differences are denoted by asterisk (*P* < 0.05*, *P* < 0.01**, *P* < 0.001***).

|  | Bacteria | Fungi |
| --- | --- | --- |
| Ag | 0.001** | 0.003** |
| Cl | 0.001** | 0.001** |
| Day | 0.001** | 0.001** |
| Ag:Cl | 0.001** | 0.636 |
| Ag:Day | 0.001** | 0.212 |
| Cl:Day | 0.001** | 0.001** |
| Ag:Cl:Day | 0.046* | 0.537 |

**Table 2.** *P*-values from multivariate GLM post-hoc results computed using mvabund highlighting difference in bacterial community composition between treatments and relative to the controls within each time point. Significant differences are denoted by asterisk (*P* < 0.05*, *P* < 0.01**, *P* < 0.001***).

| Treatment 1 |  | Treatment 2 |  | Day |  |  |  |
| --- | --- | --- | --- | --- | --- | --- | --- |
| Ag | Cl | Ag | Cl | 3 | 7 | 30 | 90 |
| (mg kg<sup>-1</sup>) | (g kg<sup>-1</sup>) | (mg kg<sup>-1</sup>) | (g kg<sup>-1</sup>) |  |  |  |  |
| 0 | 0 | 0 | 1 | <0.001*** | <0.001*** | <0.001*** | <0.001*** |
| 0 | 0 | 0 | 2 | <0.001*** | <0.001*** | <0.001*** | <0.001*** |
| 0 | 1 | 0 | 2 | 0.005** | 0.002** | 0.001** | 0.006** |
| 1 | 0 | 1 | 1 | <0.001*** | <0.001*** | <0.001*** | <0.001*** |
| 1 | 0 | 1 | 2 | <0.001*** | <0.001*** | <0.001*** | <0.001*** |
| 1 | 1 | 1 | 2 | <0.001*** | <0.001*** | 0.002** | 0.002** |
| 10 | 0 | 10 | 1 | <0.001*** | <0.001*** | <0.001*** | <0.001*** |
| 10 | 0 | 10 | 2 | <0.001*** | <0.001*** | <0.001*** | <0.001*** |
| 10 | 1 | 10 | 2 | <0.001*** | 0.001** | <0.001*** | 0.014* |
| 0 | 0 | 1 | 0 | 0.039* | <0.001*** | 0.047* | 0.185 |
| 0 | 0 | 10 | 0 | <0.001*** | <0.001*** | <0.001*** | <0.001*** |
| 1 | 0 | 10 | 0 | <0.001*** | <0.001*** | <0.001*** | 0.003** |
| 0 | 1 | 1 | 1 | 0.039* | 0.011* | 0.041* | 0.081 |
| 0 | 1 | 10 | 1 | <0.001*** | <0.001*** | <0.001*** | 0.046* |
| 1 | 1 | 10 | 1 | <0.001*** | <0.001*** | <0.001*** | 0.076 |
| 0 | 2 | 1 | 2 | <0.001*** | 0.272 | 0.041* | 0.039* |
| 0 | 2 | 10 | 2 | <0.001*** | <0.001*** | 0.002** | 0.071 |
| 1 | 2 | 10 | 2 | <0.001*** | <0.001*** | 0.019* | 0.111 |

**Fig. 1.**
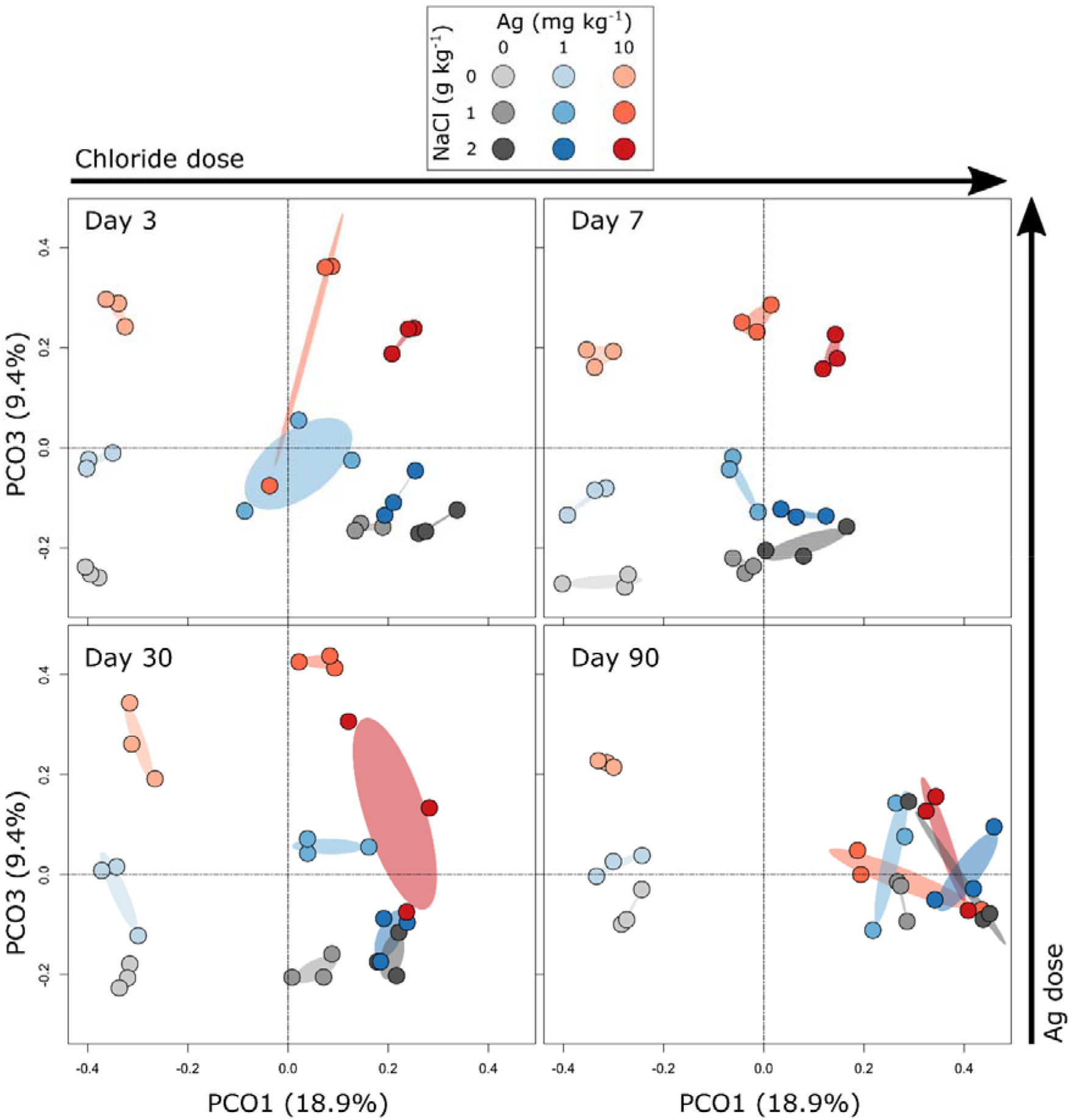
Principal coordinate analysis (PCoA) ordination illustrating differences in the composition of bacterial communities with chloride and nano-silver amendment over time. The ellipses represent standard deviations. The arrows on the right and top of the graph highlight that there was a consistent direction of change in community composition with increasing Cl and Ag dose.

The OTUs that were most strongly associated with differences in community composition between treatments were obtained from the multivariate GLMs and assessed independently using generalized linear mixed-effects models (Fig. 2). This highlighted 70 OTUs that were influenced by at least one treatment relative to the controls. Of these, 16% (11/70) showed a main effect of silver only; 34% (24/70) showed a main effect of chloride only; and 50% (35/70) showed main effects of both silver and chloride (Fig. 2). For 43% (15/35) of OTUs that showed main effect of both silver and chloride, we detected significant interactions indicating that the response to silver was influenced by soil chloride content (Fig. 2).

**Fig. 2.**
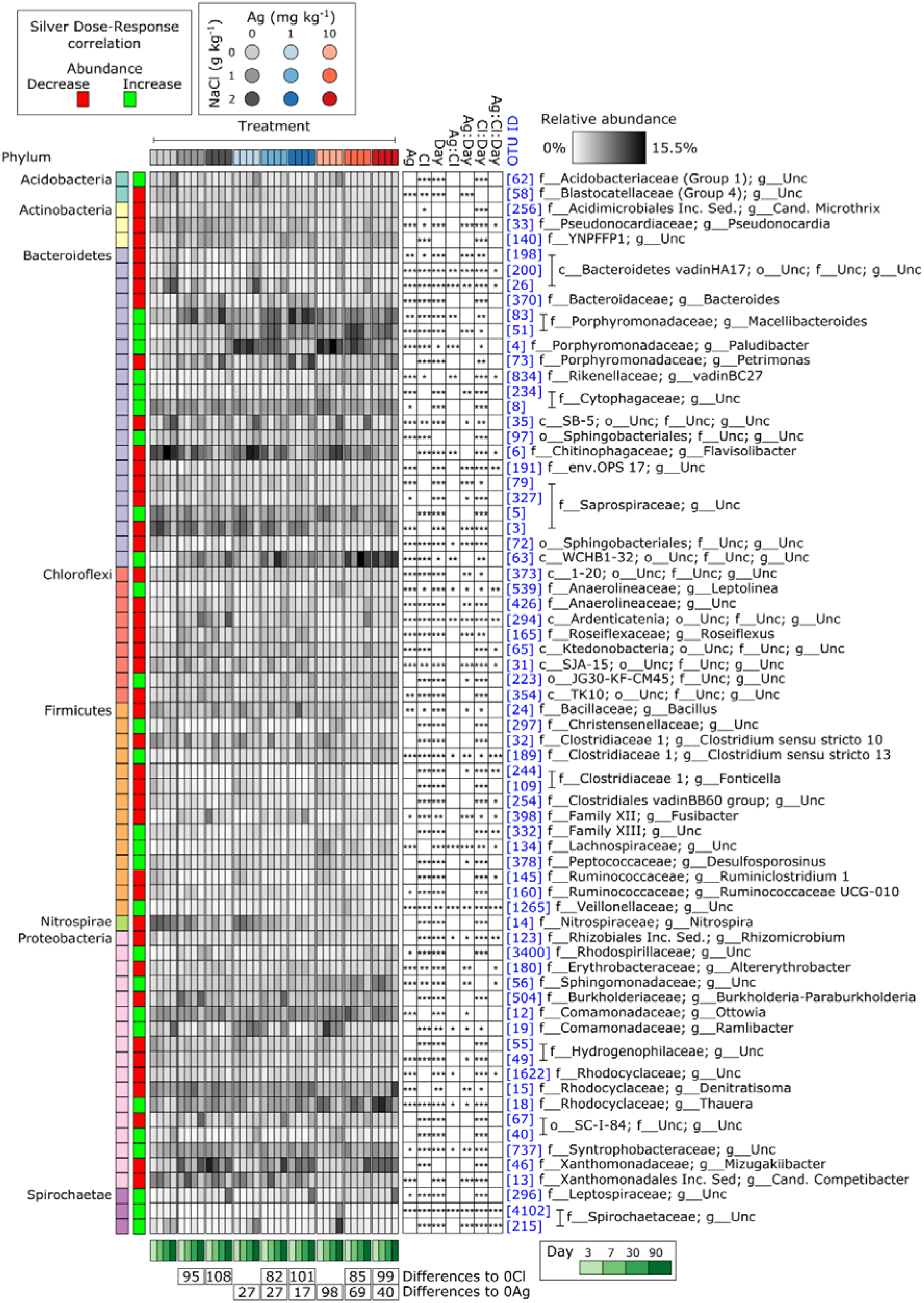
Heatmap of the relative abundances of 70 bacterial OTUs that differed significantly from respective controls in at least one treatment combination. Each column of the heatmap represents the mean relative abundance of each treatment (n = 3). The red and green boxes on the left indicate if the OTU is positively (green) or negatively (red) correlated with silver dose. The numbers below the heatmap show the total numbers of significant responses to a particular treatment relative to the respective controls. The OTU IDs are consistent throughout the manuscript. The phylum of each OTU is indicated by the colors on the left of the heatmap. The boxes on the right-hand side indicate main responses and interaction effects of each OTU.

Of the 11 OTUs that showed a main effect of silver only, seven were negatively and four were positively associated with silver dose (Fig. 2). Similarly, of the 24 OTUs that showed a main effect of chloride only, 14 were negatively and 10 were positively associated with silver dose (Fig. 2). Of the 35 OTUs that showed a main effect of both silver and chloride, 21 were negatively associated with silver dose, for which five exhibited a significant silver:chloride interaction; and 14 were positively associated with silver dose, of which ten exhibited a significant silver:chloride interaction (Fig. 2).

### 3.2. Soil fungal community composition

Fungal communities were dominated by representatives of the Ascomycota, Basidiomycota, Chytridiomycota, Mortierellomycota, Mucoromycota and Rozellomycota (Fig. S2). Fungal community composition was significantly influenced by silver and chloride dose, as well as time (Table 1, 3). Chloride content did not significantly influence the response of fungal communities to silver (Table 1, 3; Fig. 3, S3). As for bacteria, the fungal OTUs that were most strongly associated with differences in community composition between treatments were obtained from the multivariate GLMs and assessed independently using generalized linear mixed-effects models (Fig. 4). This highlighted 11 OTUs that were influenced by at least one treatment relative to the controls. Of these, 36% (4/11) of OTUs showed a main effect of silver addition only, 45% (5/11) showed a main effect of chloride only and 18% (2/11) showed a main effect of both silver and chloride (Fig. 4).

**Table 3.** *P*-values from multivariate GLM post-hoc results computed using mvabund highlighting difference in fungal community composition between treatments and relative to the controls within each time point. Significant differences are denoted by asterisk (*P* < 0.05*, *P* < 0.01**, *P* < 0.001***).

| Treatment 1 |  | Treatment 2 |  | Day |  |  |  |
| --- | --- | --- | --- | --- | --- | --- | --- |
| Ag | Cl | Ag | Cl | 3 | 7 | 30 | 90 |
| (mg kg<sup>-1</sup>) | (g kg<sup>-1</sup>) | (mg kg<sup>-1</sup>) | (g kg<sup>-1</sup>) |  |  |  |  |
| 0 | 0 | 0 | 1 | 0.014* | 0.100 | 0.083 | 0.057 |
| 0 | 0 | 0 | 2 | 0.023* | 0.016* | 0.154 | 0.321 |
| 0 | 1 | 0 | 2 | 0.249 | 0.050 | 0.256 | 0.540 |
| 1 | 0 | 1 | 1 | 0.023* | 0.035* | 0.012* | 0.032* |
| 1 | 0 | 1 | 2 | 0.021* | 0.003** | 0.039* | 0.041* |
| 1 | 1 | 1 | 2 | 0.231 | 0.003** | 0.021* | 0.231 |
| 10 | 0 | 10 | 1 | 0.036* | 0.009** | 0.003** | 0.021* |
| 10 | 0 | 10 | 2 | 0.031* | 0.002** | 0.002** | 0.021* |
| 10 | 1 | 10 | 2 | 0.118 | 0.003** | 0.384 | 0.048* |
| 0 | 0 | 1 | 0 | 0.012* | 0.118 | 0.321 | 0.601 |
| 0 | 0 | 10 | 0 | 0.046* | 0.008** | 0.003** | 0.825 |
| 1 | 0 | 10 | 0 | 0.016* | 0.020* | 0.011* | 0.514 |
| 0 | 1 | 1 | 1 | 0.231 | 0.560 | 0.176 | 0.061 |
| 0 | 1 | 10 | 1 | 0.184 | 0.023* | 0.278 | 0.082 |
| 1 | 1 | 10 | 1 | 0.084 | 0.027* | 0.030* | 0.143 |
| 0 | 2 | 1 | 2 | 0.258 | 0.202 | 0.478 | 0.666 |
| 0 | 2 | 10 | 2 | 0.740 | 0.003** | 0.321 | 0.391 |
| 1 | 2 | 10 | 2 | 0.431 | 0.003** | 0.335 | 0.143 |

**Fig. 3.**
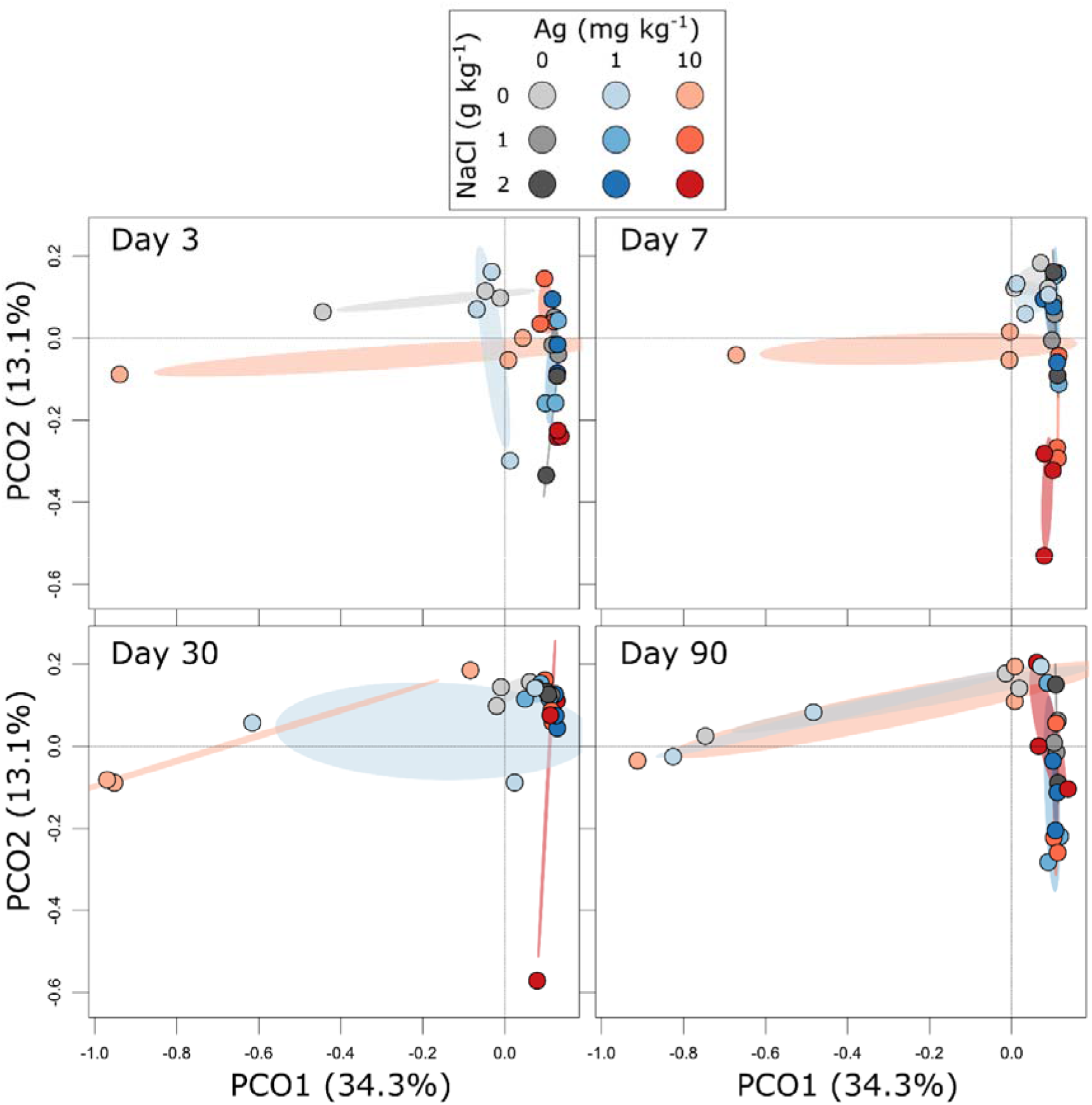
Principal coordinate analysis (PCoA) ordination illustrating differences in the composition of fungal communities with chloride and nano-silver amendment over time. The ellipses represent standard deviations.

**Fig. 4.**
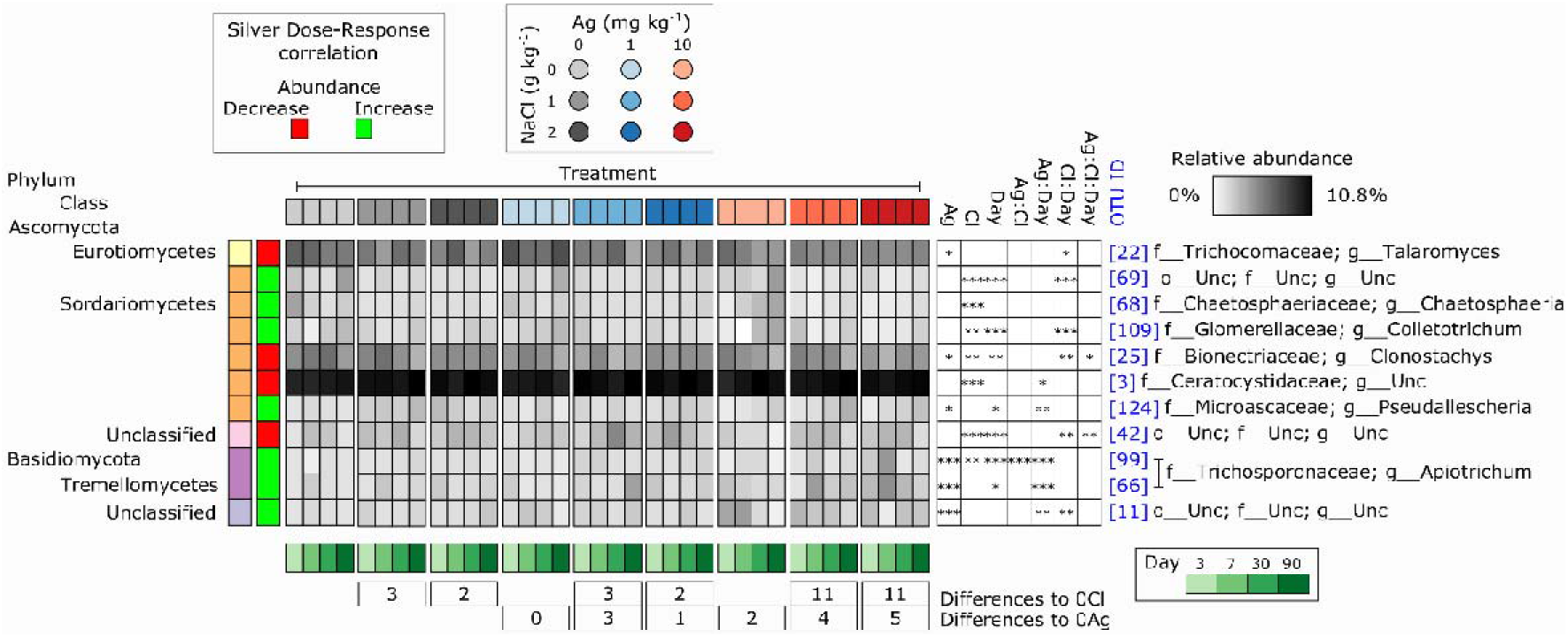
Heatmap of the relative abundances of 11 fungal OTUs that differed significantly from respective controls in at least one treatment combination. Each column of the heatmap represents the mean relative abundance of each treatment (n = 3). The red and green boxes on the left indicate if the OTU is positively (green) or negatively (red) correlated with silver dose. The numbers below the heatmap show the total numbers of significant responses to a particular treatment relative to the respective controls. The OTU IDs are consistent throughout the manuscript. The phylum of each OTU is indicated by the colors on the left of the heatmap. The boxes on the right-hand side indicate main responses and interaction effects of each OTU.

Of the four OTUs that showed a main effect of silver only, one was negatively and three were positively associated with silver dose (Fig. 2). Of the five taxa that that showed a main effect of chloride only, two were negatively and three were positively associated with silver dose (Fig. 2). Of the remaining two taxa that showed a main effect of both silver and chloride, one was positively and another was negatively associated with silver dose. While the main effect responses to silver for Ascomycota were mixed, the three Basidiomycota OTUs were all positively correlated with increasing silver dose (Fig. 4).

### 3.3. Alpha diversity of soil microbial communities

Relative to the controls, the alpha diversity of bacterial communities was significantly affected only by silver addition (Table 4, Fig. S4). The effects of silver were apparent for all alpha diversity metrics, were independent of chloride and time, and more apparent three days after treatment (Table 4). For fungi, silver and chloride did not significantly influence any of the alpha diversity metrics (Table 4).

**Table 4.**
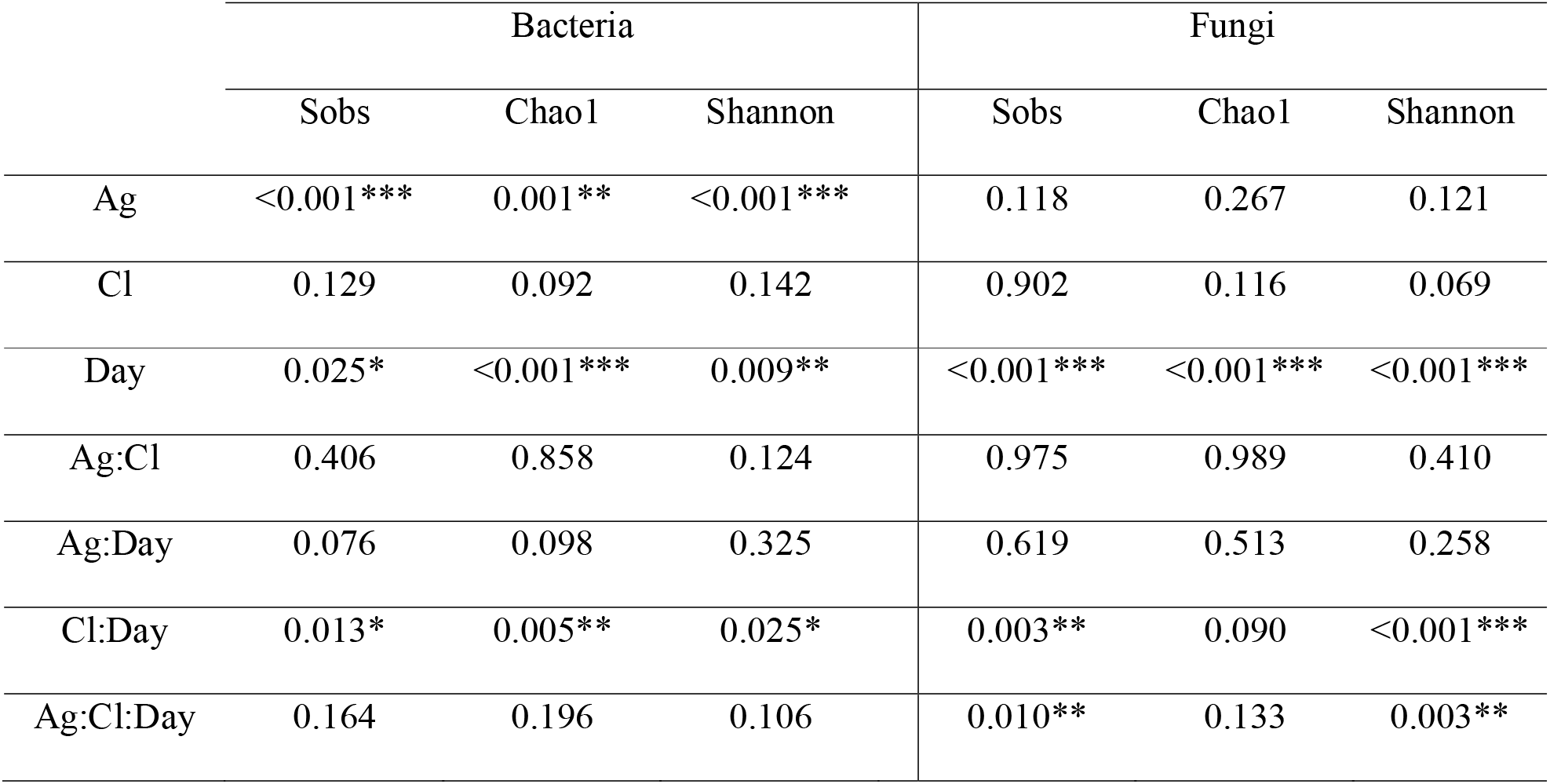
*P*-values from linear mixed-effects models computed using lme4 and lmerTest highlighting the main and interaction effects of silver, chloride and day on three measures of microbial diversity. Significant differences are denoted by asterisk (*P* < 0.05*, *P* < 0.01**, *P* < 0.001***).

|  | Bacteria |  |  | Fungi |  |  |
| --- | --- | --- | --- | --- | --- | --- |
|  | Sobs | Chao1 | Shannon | Sobs | Chao1 | Shannon |
| Ag | <0.001*** | 0.001** | <0.001*** | 0.118 | 0.267 | 0.121 |
| Cl | 0.129 | 0.092 | 0.142 | 0.902 | 0.116 | 0.069 |
| Day | 0.025* | <0.001*** | 0.009** | <0.001*** | <0.001*** | <0.001*** |
| Ag:Cl | 0.406 | 0.858 | 0.124 | 0.975 | 0.989 | 0.410 |
| Ag:Day | 0.076 | 0.098 | 0.325 | 0.619 | 0.513 | 0.258 |
| Cl:Day | 0.013* | 0.005** | 0.025* | 0.003** | 0.090 | <0.001*** |
| Ag:Cl:Day | 0.164 | 0.196 | 0.106 | 0.010** | 0.133 | 0.003** |

## 4. Discussion

In this study we tested the hypotheses that: 1) metallic Ag-NPs entering soil via wastewater treatment processing influence the diversity of bacterial and fungal communities, and that 2) these effects would be more apparent in soils with elevated chloride content, as we have previously demonstrated, using samples from the same experiment, that chloride addition increases silver bioavailability (Wang et al., 2016).

With respect to the first hypothesis, our findings indicate that wastewater treatment processed Ag-NPs influence the composition of bacterial and fungal communities. However, wastewater treatment processed Ag-NPs only appeared to influence the alpha diversity of bacterial communities, suggesting that they are more sensitive to the Ag treatments than fungi. These findings are important because most silver enters soils in biosolids and is present as Ag_2_S, which previous studies have indicated to be less toxic to microbes and other organisms (Judy et al., 2015; Levard et al., 2013; Reinsch et al., 2012; Wang et al., 2018a). Hence, despite reductions in the toxicity of Ag-NPs during wastewater treatment processing, they may still affect soil microbial diversity, even at environmentally realistic concentrations. Previously, wastewater treatment processed Ag-NPs have been shown to adversely affect ammonia oxidizing bacteria (Kraas et al., 2017; Schlich et al., 2018). Hence our findings greatly expand the range of taxa affected by wastewater treatment processed Ag-NPs.

Contrary to the results of Judy et al. (2015), our study also indicates a positive association between silver dose and the magnitude of change in bacterial community composition (Fig. 1). This may be due to an increase in the abundance of Ag^+^ ions at larger Ag-NPs loading rates. For example, sulfidation of Ag-NPs during aging processes can result in a shell-core structure, whereby the outer layers of Ag-NPs are sulfidized while the core remains pristine (Elechiguerra et al., 2005; Kaegi et al., 2011). Consequently, there may be a reservoir of metallic Ag^+^ available for release, which elicits effects on soil microbial communities until reserves are exhausted and sulfidation of this core is complete.

With respect to the second hypothesis, our study demonstrated that silver-associated changes in bacterial community composition were affected by soil chloride content, with more acute responses to silver being observed in soils with elevated chloride content (Fig. 1). In contrast, chloride did not influence silver-associated changes in fungal community composition or the alpha diversity of bacterial or fungal communities. Interestingly, the effect of silver dose persisted until day 90 in samples with no chloride addition, but was diminished in all chloride amended treatments. As chloride has been shown to increase the bioavailability and solubility of silver in soil (Wang et al., 2016), it is possible that in higher chloride treatments the Ag_2_S shell would have been lost more rapidly along with the Ag^+^ core. Consequently, in the elevated chloride treatments, the influence of wastewater treatment processed Ag-NPs on microbes may have been more acute because Ag^+^ was dispersed and converted to less toxic Ag_2_S more rapidly. This is supported by our previous work, which demonstrated that after 90 days, metallic Ag^+^ was detectable in no-chloride treatments, but not in elevated chloride treatments (Wang et al., 2016).

The majority (66%) of affected bacterial taxa showed a main effect of silver addition and many of these responses were significantly altered by chloride. Most bacterial taxa responding to treatments were negatively correlated with increasing silver dose. Interestingly, OTU 378 (*Desulfosporosinus*), which is closely related to bacteria known to reduce sulfate, was positively correlated with the addition of wastewater treatment processed Ag-NPs. This may be the result of a greater availability of sulfate groups due to the addition of Ag_2_S. Relative to bacterial taxa, the responses of individual fungal OTUs were less numerous. Over half of the taxa affected by treatments exhibited a significant main effect of silver addition, and of these, the majority increased in relative abundance independently of chloride level. In general, fungi appeared to be less sensitive to treatments than bacteria, with fewer OTUs individually responding to treatments and no effect of silver or chloride being apparent on overall fungal diversity.

In summary, not only did wastewater treatment processed Ag-NPs significantly alter bacterial community composition, the nature of this effect on the whole community, as well as on individual taxa, was affected by soil chloride concentration. While the silver treatments also significantly affected fungal community composition, these effects were independent of chloride dose and only a small number of taxa were found to respond. Additionally, while bacterial alpha diversity was affected by silver addition, the alpha diversity of fungal communities was not affected by any of the treatments. These factors suggest that fungal communities may be less susceptible to wastewater treatment processed Ag-NPs than bacteria. This is supported by previous, culture based, studies which found that fungi were unaffected by silver doses as high as 25 mg kg^−1^ in some instances (Kim et al., 2012) whereas bacterial cultures were consistently affected in various studies even at low dose (Choi and Hu, 2008; Dhas et al., 2014; Gajjar et al., 2009; Kim et al., 2007; Radzig et al., 2013).

## Conclusions

Given that Ag-NPs are widely used and are being released into the environment (Vance et al., 2015), there is an urgent need to understand their impacts, including those on soil microbial communities. Here, we demonstrated that wastewater treatment processed Ag-NPs can affect soil bacterial and fungal community composition, and that for bacterial communities these effects are modulated by soil chloride concentration. We demonstrated a dose effect on bacterial community composition by wastewater treatment processed Ag-NPs, as well as the decrease of this dose effect relationship over time in the presence of increased soil chloride levels. This highlights that even realistic applications of Ag-NPs after wastewater treatment can significantly influence soil microbial community composition and that these effects may differ depending on soil salinity.

## Acknowledgements

The authors gratefully acknowledge financial support from The University of Queensland for an Early Career Researcher Award to PGD. CF gratefully acknowledges funding from the Australian Government’s Department of Education and Training in the form of an Australian Government Research Training Program Scholarship administered by The University of Queensland. PMK is the recipient of an Australian Research Council (ARC) Future Fellowship (ARC FT120100277).

