## Supplementary Information for "Soil chloride content influences the response of bacterial but not fungal diversity to silver nanoparticles entering soil via wastewater treatment processing"

Contents:

**Fig. S1** Relative abundance of bacterial phyla.........................................................................2

**Fig. S2** Relative abundance of fungal phyla………………………………………………...3

**Fig. S3** PCoA of fungal samples, better separation of samples clumped on Fig. 3……………4

**Fig. S4** The numbers of observed OTUs in bacterial and fungal samples over time……..…...5

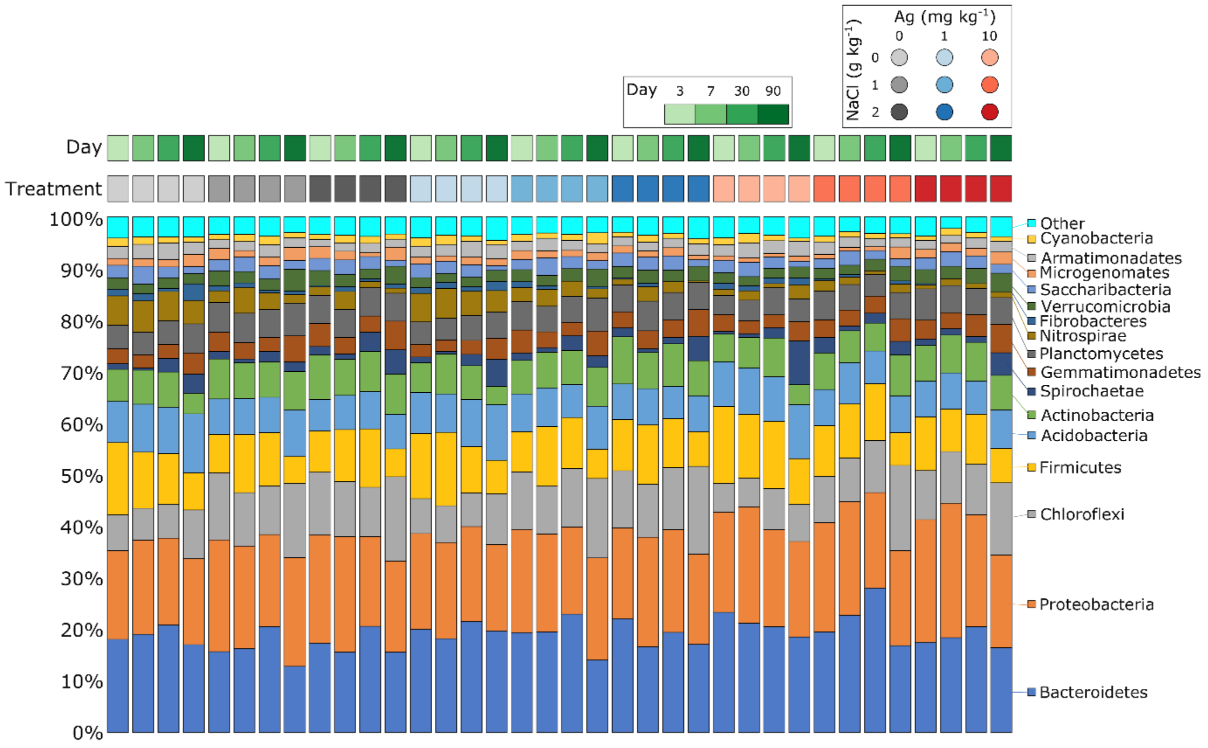


**Fig. S1** The relative abundances of bacteria phyla in control, and Cl and Ag amended soils over time. All phyla representing <1% relative abundance are combined as "Other".

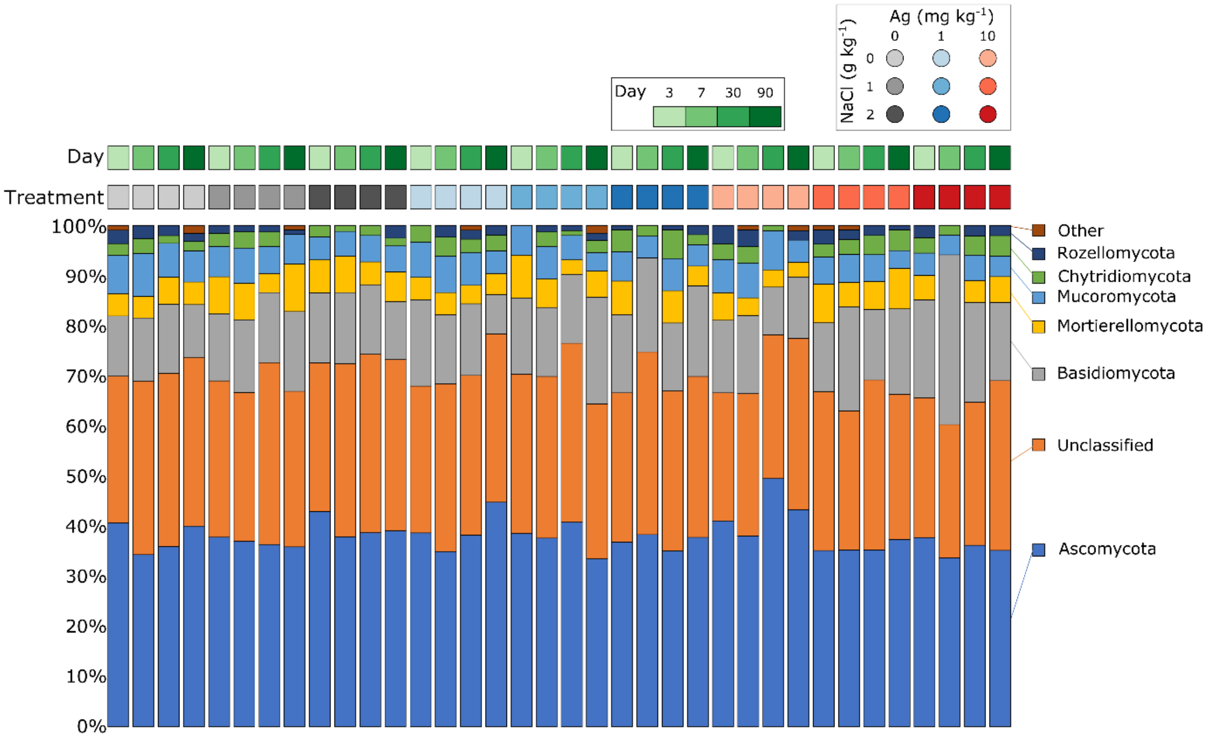


**Fig. S2** The relative abundances of fungal phyla in control, and Cl and Ag amended soils over time. All phyla representing <1% relative abundance overall are combined as "Other".

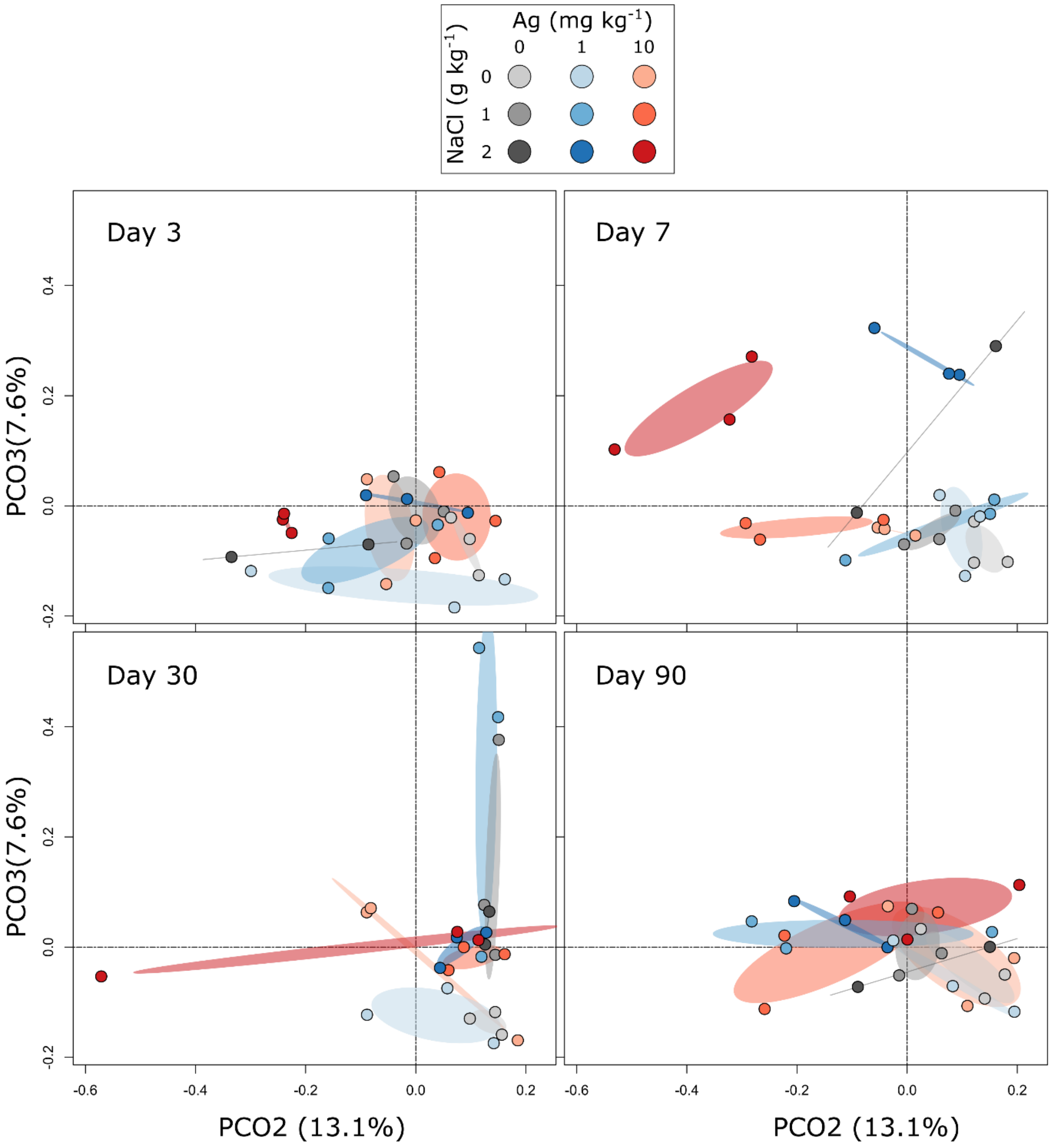


**Fig. S3** Principal coordinate analysis (PCoA) ordination summarizing variation in the composition of fungal communities associated with soils amended with different doses of chloride and/or Ag over time. The ellipses represent standard deviations. This PCoA provides better visual separation of samples clumped along PCO1 in Fig. 3.

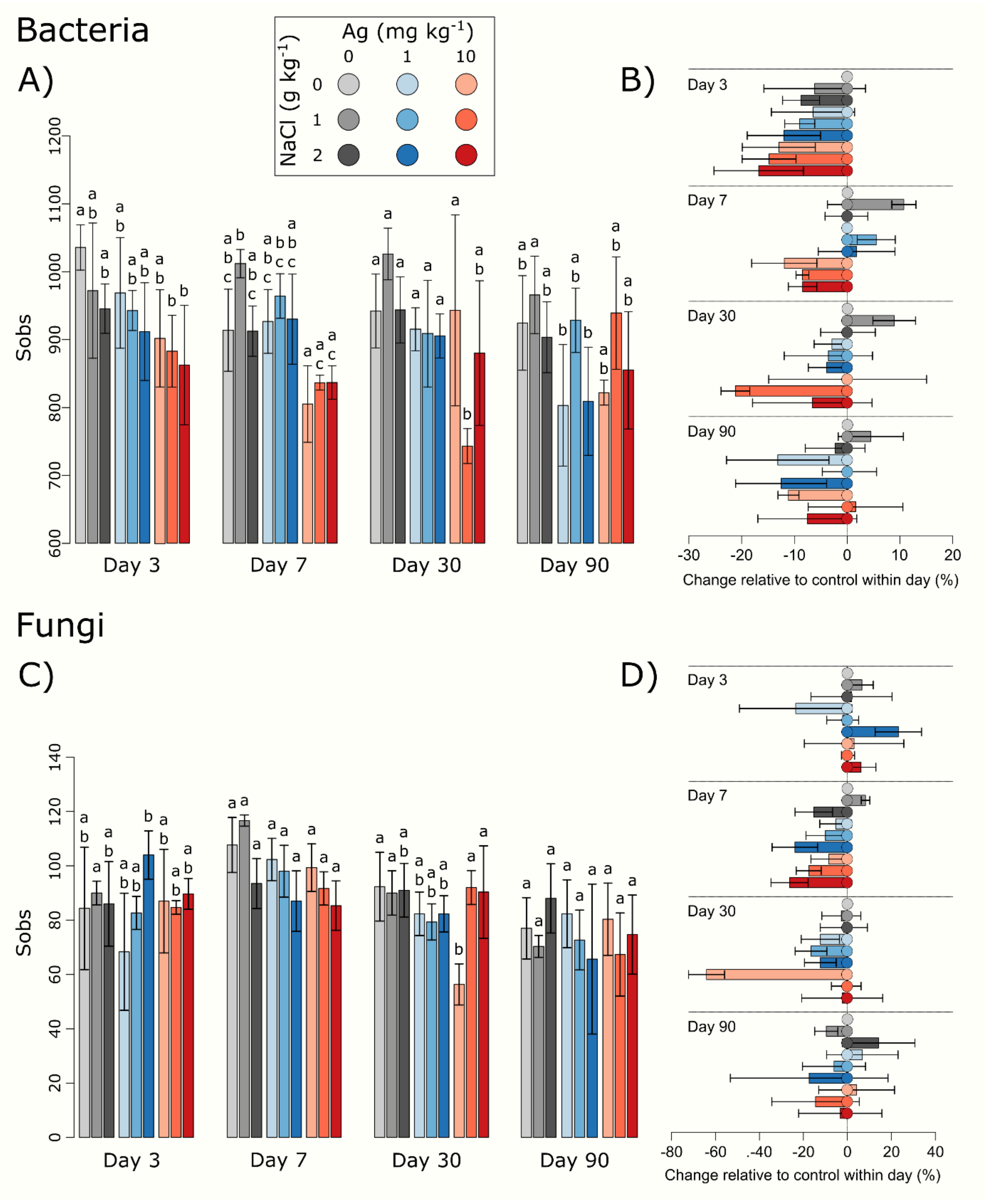


**Fig. S4** The numbers of observed (Sobs) bacterial (A) and fungal (C) OTUs over time. Letters indicate significant differences within days only. Error bars represent standard deviations. Significant differences are denoted by letter groups. Percentage change from the no treatment control is shown by the horizontal bars for both bacterial (B) and fungal (D) OTUs
